# Astrocyte calcium dysfunction causes early network hyperactivity in Alzheimer’s Disease

**DOI:** 10.1101/2022.04.26.489446

**Authors:** Disha Shah, Willy Gsell, Jérôme Wahis, Emma S. Luckett, Tarik Jamoulle, Ben Vermaercke, Pranav Preman, Daan Moechars, Véronique Hendrickx, Tom Jaspers, Katleen Craessaerts, Katrien Horré, Leen Wolfs, Mark Fiers, Matthew Holt, Dietmar Rudolf Thal, Zsuzsanna Callaerts-Vegh, Rudi D’Hooge, Rik Vandenberghe, Uwe Himmelreich, Vincent Bonin, Bart De Strooper

## Abstract

Dysfunctions of network activity and functional connectivity (FC) represent early events in Alzheimer’s disease (AD), but the underlying mechanisms remain unclear. Astrocytes regulate neuronal activity in the healthy brain, but their involvement in early network hyperactivity in AD is unknown. We show increased FC in the human cingulate cortex, several years before amyloid deposition. We found the same early cingulate FC disruption and neuronal hyperactivity in *App^NL-F^* mice. Crucially, these network disruptions are accompanied by decreased astrocyte calcium signaling. Recovery of astroglial calcium activity normalizes neuronal hyperactivity and FC, as well as seizure susceptibility and day/night behavioral hyperactivity. In conclusion, we show for the first time that astrocytes mediate initial features of AD and drive clinically relevant phenotypes.

## Main text

Neuronal deficits contribute to network disruptions that characterize the initial stages of Alzheimer’s disease (AD). Mouse models of amyloid pathology show early neuronal hyperactivity, which manifests as increased calcium signaling in neurons (1, 2). Neuronal deficits can also lead to alterations of functional connectivity (FC), measured with resting-state functional Magnetic Resonance Imaging (rsfMRI), a translational tool that is used extensively in clinical research. Indeed, increased FC of cortical and hippocampal regions occurs before amyloid plaque deposition in *App* mouse models (3–5) and in individuals at risk for AD, such as APOE-ε4 carriers (6). Network activity and FC thus represent clinically relevant readouts for early AD, and defining the mechanisms that underlie these changes can be important for therapeutic development.

Disruption of neuronal activity could be mediated by functional deficits of astrocytes. When amyloid plaques develop, astrocytes become reactive and, together with microglia, contribute to neuro-inflammatory processes (7). However, astrocytes are also implicated in maintaining homeostasis of neuronal circuits (8,9) and regulate neuronal activity through, among others, fluctuations of their intracellular calcium signals (10) and subsequent secretion of gliotransmitters (11,12). It is known that reactive astrocytes produce increased spontaneous calcium oscillations in several neurological disorders (13), and near amyloid plaques (14). It is unclear, however, if astrocytes already show functional deficits before amyloid plaque deposition. An early disruption of astrocyte calcium signaling could impede their ability to regulate neuronal activity, which in turn could possibly instigate or sustain network hyperactivity in AD.

We aimed at identifying early responding brain regions that show FC disruptions in cognitively intact human individuals and the *App^NL-F^* mouse model (15), prior to amyloid plaque deposition. We then zoomed in on in-vivo functional changes at the cellular level in neurons and astrocytes. Finally, we modulate astrocyte activity to prove their involvement in neuronal disruptions, FC deficits and clinically important behavior readouts. Our results demonstrate a fundamental contribution of astrocytes to initial features of AD.

## Results

### Increased cingulate functional connectivity prior to plaque deposition in humans and mice

RsfMRI allows non-invasive whole brain measurements of FC, using the blood-oxygen-level-dependent (BOLD) contrast as an indirect measure for brain activity. We analyzed rsfMRI data of cognitively intact participants from the Flemish Prevent AD Cohort KU Leuven (F-PACK)(16), who were scanned with amyloid-PET at two time points (7±0.3 years between two scans). We divided the group into individuals who either show no change in amyloid load over time (controls, ‘Centiloid’ CL at baseline scan 7.1±1.1, CL at follow-up scan 6.1.0±1.6, N=24, p=0.83) or increasing amyloid load (amyloid accumulators, CL at baseline 7.7±2.1, CL at follow-up scan 42±5.7, N=10, p<0.0001) (**Fig. 1A**). We applied a threshold of CL=23.5 to discriminate amyloid accumulators and controls (16,17). We compared the rsfMRI scans acquired at baseline between these groups to assess whether there are changes in BOLD FC prior to longitudinal increases in amyloid load. The mean FC matrices (**Fig. 1B**) show increased anterior-posterior cingulate FC in amyloid accumulators (False Discovery Rate, FDR corrected at p<0.05, p=0.0005), and we found a significant difference using the change in amyloid as a covariate for this connection (FDR corrected, p=0.006). This was confirmed by the mean FC maps (FDR corrected, p<0.05) and statistical difference map (uncorrected, p<0.001) of the anterior cingulate cortex, which show increased anterior-posterior FC in amyloid accumulators (**Fig. 1C**). We used the individual FC maps to quantify anterior-posterior FC of the cingulate cortex (**Fig. 1D**) (z-scores for control group 0.25 ± 0.025 for N=24 individuals, amyloid accumulators 0.54 ± 0.083 for N=10 individuals, p=0.0001). Moreover, we found a positive correlation between the increase in amyloid load and baseline anterior-posterior cingulate FC in the amyloid accumulators (r=0.86, p=0.0016) (**Fig. 1E**), suggesting that cingulate FC has predictive value for downstream amyloid accumulation.

**Fig 1:**
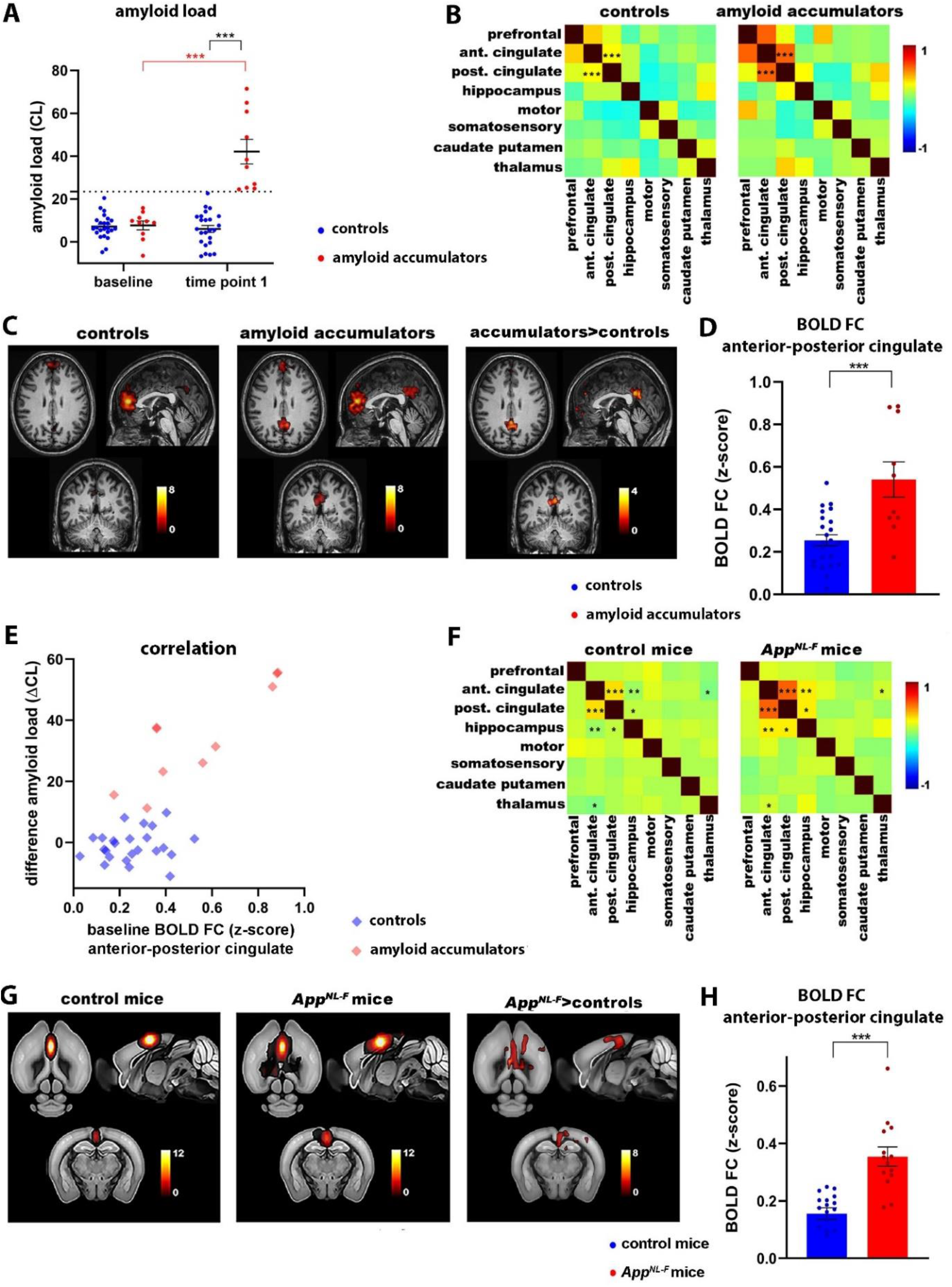
Increased FC of the anterior cingulate cortex in the human brain and *App^NL-F^* mice. **A)** Amyloid load (CL values±SEM) for controls (N=24) and amyloid accumulators (N=10). Threshold at CL=23.5 is indicated by the dotted line. ***p<0.001, two-way repeated ANOVA with Sidak correction. **B)** Mean FC matrices of controls (N=24) and amyloid accumulators (N=10). ***p<0.001, two-sample T-test, FDR corrected. Color scale shows z-scores. **C)** Mean BOLD FC map of the anterior cingulate cortex of controls and amyloid accumulators (FDR corrected, p<0.05). Color scale shows T-values, i.e. FC between the anterior cingulate cortex and all other voxels in the brain. Statistical map shows anterior cingulate connections that are increased in amyloid accumulators versus controls, two-sample T-test, uncorrected, p<0.001. **D)** BOLD FC (z-scores ±SEM) between the anterior and posterior cingulate cortex. ***p<0.001, two-sample T-test. **E)** Correlation between change in amyloid load (Δ CL between baseline and time point 1) and BOLD FC (between the anterior and posterior cingulate cortex) at baseline for controls (r=0.14, p=0.52) and amyloid accumulators (r=0.86, p=0.0016). **F)** Mean FC matrices of 3 months old control (N=17) and *App^NL-F^* mice (N=14). *p<0.05, **p<0.01, ***p<0.001, two-sample T-test, FDR corrected. Color scale shows z-scores. **G)** Mean BOLD FC map of the anterior cingulate cortex of control and *App^NL-F^* mice (FDR corrected, p<0.05). Color scale shows T-values. Statistical map shows anterior cingulate connections that are increased in *App^NL-F^* mice versus controls, two-sample T-test, FDR corrected, p<0.05. **H)** BOLD FC (z-scores ±SEM) between the anterior and posterior cingulate cortex. ***p<0.001, two-sample T-test.

Although increased FC of cortical and hippocampal networks has been demonstrated in patients who have amyloid plaques and are at risk for developing AD (6), we show here that cognitively normal individuals show early increases of cingulate BOLD FC, several years before reaching the threshold for amyloid positivity on PET scans. This indicates that increased FC of the cingulate circuitry represents a very early functional symptom of incipient AD.

We next assessed whether similar early FC changes can be observed in the (preclinical) early disease stages, i.e. at 3 months of age, in the *App^NL-F^* mouse model of AD. The cingulate cortex and hippocampus show increased FC before amyloid plaque deposition. FC between the anterior and posterior cingulate cortex shows the most significant change (FDR corrected, anterior-posterior cingulate cortex p=0.0004) (**Fig. 1F**). The mean FC maps and statistical difference map of the anterior cingulate cortex confirm increased anterior-posterior FC in *App^NL-F^* mice compared to *App^NL^* controls (FDR corrected, p<0.05) (**Fig. 1G**). We used the individual maps to quantify FC between the anterior and posterior cingulate cortex (**Fig. 1H**) (z-scores for control mice 0.16 ± 0.020 for N=17 mice, *App^NL-F^* 0.35 ± 0.033 for N=14 mice, p<0.0001). Thus, the *App^NL-F^* mouse model shows similar early increases of BOLD FC in the cingulate cortex as humans, emphasizing the clinical relevance of the mouse model and the importance of the cingulate cortex during early disease stages.

### Astrocytes regulate network functional connectivity in the healthy brain

While we know that astrocytes support the regulation of local neuronal activity (11, 12), it remains unknown whether astrocytes are also involved in regulation of network connectivity. We used wild-type C57BL/6 mice to assess whether modulation of intracellular calcium signals in astrocytes can modify FC in the healthy brain.

We used adeno-associated viruses (AAV) to express different transgenes in astrocytes under the GFAP promoter. First, we expressed hM3Dq Designer Receptors Exclusively Activated by Designer Drugs (DREADDs) (18), which respond to systemic injection of clozapine-N-oxide (CNO) by calcium release from the endoplasmatic reticulum (ER) through the inositol 1,4,5-trisphosphate (IP3) pathway(18). Conversely, we expressed the Pleckstrin Homology (PH) domain of Phospholipase C (PLC)-like protein p130 (p130PH), which buffers IP3 and impedes calcium release (19). We stereotactically injected all AAVs in the cingulate cortex and performed several control experiments to demonstrate the specificity and efficiency of this strategy. **Fig. S1** shows the AAV injection site, and specific expression of AAVs in astrocytes. Furthermore, co-injection of AAVs targeted to neurons (hSYN promoter, eGFP) or astrocytes (GFAP promoter, mCherry) showed no co-expression of the two fluorophores, validating the cell type specificity of our strategy.

Next, we used in-vivo two-photon calcium imaging of astrocytes expressing GCaMP6f to confirm that activation of DREADDs through systemic injection of CNO increases the amplitude and frequency of astrocyte calcium signals, while p130PH expression decreases these parameters in wild-type C57BL/6 mice (**Fig. S2**).

We then show that modulating astrocyte calcium signals alters FC of the cingulate cortex as measured with wide field calcium imaging of neurons, after intravenous administration of AAV-PHP.eB under the hSYN promoter to enable widespread expression of Gcamp6f in neurons (**Fig. S3**). Furthermore, modifying astrocyte calcium activity also affects BOLD FC measured with rsfMRI (**Fig. S4).** These data demonstrate an important role of astrocytes in controlling FC and supporting large-scale functional organization of the healthy brain, prompting us to test if astrocytes calcium signaling is altered in *App^NL-F^* mice.

### Astrocytes show decreased calcium signaling at early stages of amyloid pathology

**I**n-vivo two-photon calcium imaging of GCaMP6f expressing astrocytes shows a decrease of astrocyte calcium signaling in *App^NL-F^* mice compared to control mice, as illustrated by the representative fluorescence calcium (ΔF/F_0_) time traces (**Fig. 2A**) and heat maps (**Fig. 2B**). The percentage of active cells is decreased in *App^NL-F^* mice (controls mCherry 64%, N=223 cells from 5 mice; *App^NL-F^* mCherry 28%, N=291 cells from 4 mice) (**Fig. 2C**). Furthermore, the frequency (number of peaks per minute for controls mCherry 0.12±0.011; *App^NL-F^* mCherry 0.04 ± 0.006, p<0.0001) and ΔF/F0 amplitude (controls mCherry 1.32±0.0133; *App^NL-F^* mCherry 1.21± 0.00763, p<0.0001) of the calcium time traces are decreased in *App^NL-F^* mice (**Fig. 2D-E**).

**Fig. 2:**
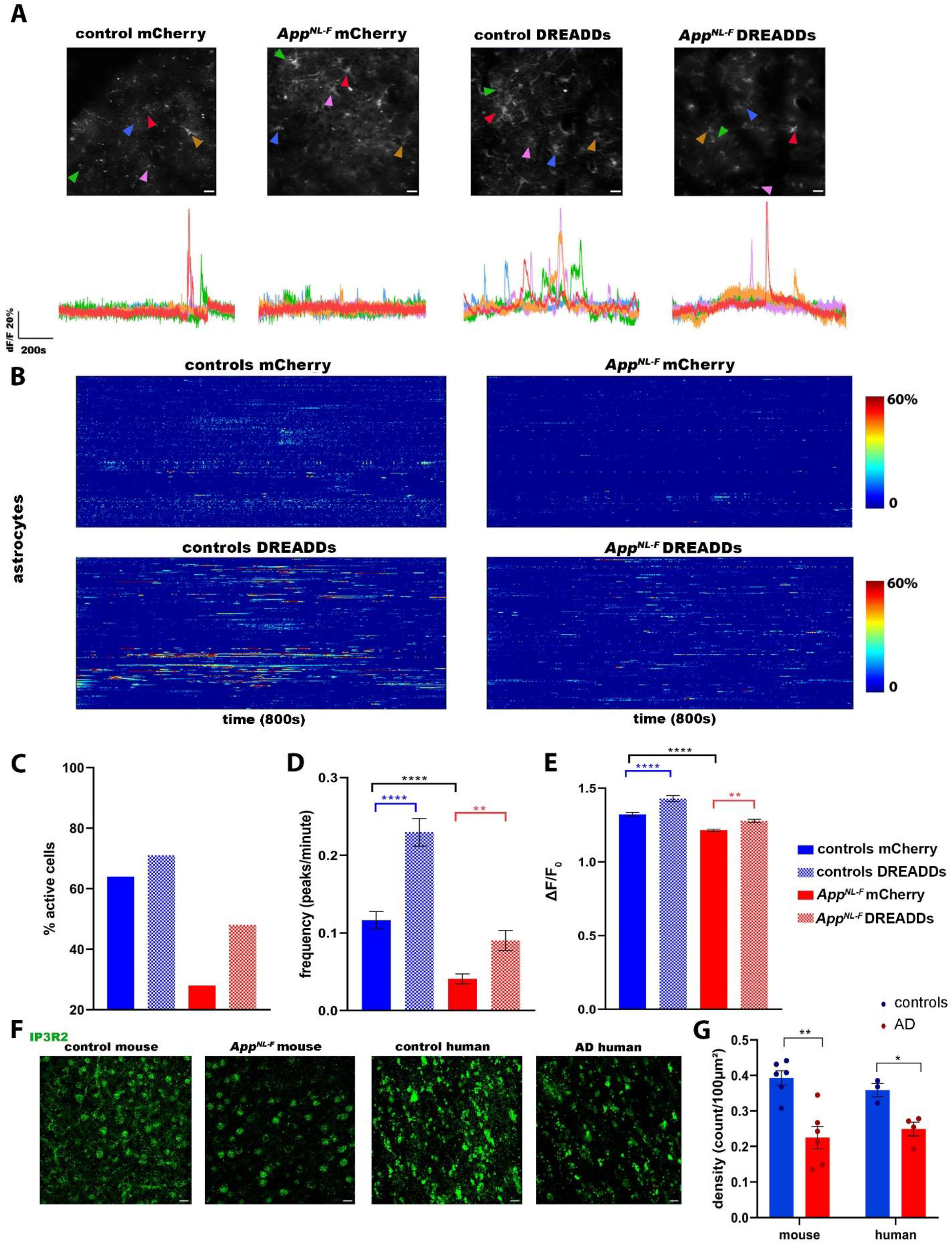
Astrocytes show decreased calcium signaling at early stages of amyloid pathology. **A)** Representative ΔF/F_0_ calcium traces of astrocytes in the cingulate cortex of 3 months old control and *App^NL-F^* mice expressing the calcium reporter GCaMP6f in addition to either control AAV (mCherry) or DREADDs in astrocytes (GFAP promoter). Scale bar=20μm. **B)** Heat maps show calcium activity (>20% of baseline, Y-axis) over time (800s, X-axis), quantifying astrocyte calcium signals in control mice expressing mCherry (N=5 mice, 223 cells) or DREADDs (N=4 mice, 263 cells), and *App^NL-F^* mice expressing mCherry (N=4 mice, 291 cells) or DREADDs (N=4 mice, 236 cells). **C-D-E)** Graphs show percentage of active cells, frequency (number of peaks per minute ±SEM) and signal amplitude (ΔF/F0 ±SEM). **p<0.01, ****p<0.0001, one-way ANOVA with Sidak correction. **F-G**) Representative images and quantification (density/100μ±SEM) of immunofluorescent staining of the IP3R2 receptor in mouse (N=6 controls, N=6 *App^NL-F^* mice) and human AD cortical tissue (N=3 controls, N=4 AD patients). Scale bar=25μm. *p<0.05, **p<0.01, two-sample T-test.

Cytosolic calcium disruptions in *App^NL-F^* mice could be caused by a deficit of calcium release from internal stores. To increase calcium signaling in astrocytes we expressed either control virus (mCherry) or DREADDs in astrocytes.

Activation of DREADDs (CNO 3mg/kg) increases the percentage of active cells (controls DREADDs 71%, N=263 cells from 4 mice; *App^NL-F^* DREADDs 48%, N=236 cells from 4 mice) (**Fig. 2C**). The frequency (controls DREADDs 0.23 ± 0.018; *App^NL-F^* DREADDs 0.09 ± 0.01; controls mCherry versus DREADDs p<0.0001; *App^NL-F^* mCherry versus DREADDs p=0.005) and ΔF/F0 amplitude (controls DREADDs 1.43 ± 0.0207; *App^NL-F^* DREADDs 1.28 ± 0.0107; controls mCherry versus DREADDs p<0.0001; *App^NL-F^* mCherry versus DREADDs p=0.001) are also increased (**Fig. 2D-E**).

However, activation of DREADDs elicits a weaker increase of calcium signaling in *App^NL-F^* mice than in controls (frequency and amplitude for controls DREADDs versus *App^NL-F^* DREADDs, p<0.0001), which could point to a downregulation of IP3 receptor type 2 (IP3R2), an intracellular calcium release channel located in astrocytes (20, 21). Indeed, immunohistochemistry analyses of the IP3R2 receptors demonstrate ~42% decreased expression in *App^NL-F^* mice (puncta density per 100μ^2^ in control mice 0.39 ± 0.020 from N=6 mice, *App^NL-F^* mice 0.23±0.032 from N=6 mice, p=0.001). Moreover, we analyzed cortical tissue from control individuals (CERAD score 0, N=3) and AD patients (CERAD score 3, N=4) and found ~31% decrease of IP3R2 expression in human AD brains (control 0.36 ±0.018, AD 0.25±0.019, p=0.01) (**Fig. 2 F-G**). Thus, astrocytes show early deficits in AD, reflected as decreased calcium signaling in *App^NL-F^* mice and decreased IP3R2 expression in *App^NL-F^* mice and human AD.

### Restoring astrocyte calcium signaling dampens early neuronal hyperactivity

We next wondered whether DREADDs-mediated modulation of astrocytes would affect neuronal activity. In line with calcium imaging results in other amyloid mouse models (22), we show that GCaMP6f-expressing cingulate neurons display hyperactive calcium signaling in 3 months old *App^NL-F^* mice, as shown in the representative calcium traces and heat maps (**Fig. 3A-B**). We found an increase in the frequency (controls mCherry 14.0 ± 0.149, N=463 neurons from N=5 mice; *App^NL-F^* mCherry 15.5 ± 0.129, N=652 neurons from N=5 mice; p<0.0001) and ΔF/F_0_ amplitude (controls mCherry 1.34±0.0168; *App^NL-F^* mCherry 1.61 ± 0.0257; p<0.0001) of calcium time traces. Activating DREADDs in astrocytes of control mice causes an increase in neuronal calcium activity, i.e. signal frequency (controls DREADDs 17.5 ± 0.232, N=640 neurons from 5 mice, p<0.0001) and ΔF/F_0_ amplitude (controls DREADDs 1.67 ± 0.0285, p<0.0001). On the other hand, recovering the functional deficit of astrocytes in *App^NL-F^* mice dampens neuronal hyperactivity, i.e. frequency *(App^NL-F^* DREADDs 14.5 ± 0.241, N=466 neurons from 5 mice, p=0.0006) and ΔF/F0 amplitude of the calcium traces *(App^NL-F^* DREADDs 1.27 ± 0.0221, p<0.0001) (**Fig. 3C-D**). Thus, restoring early astrocyte disruptions in *App^NL-F^* mice decreases neuronal activity to levels comparable to control mice.

**Fig. 3:**
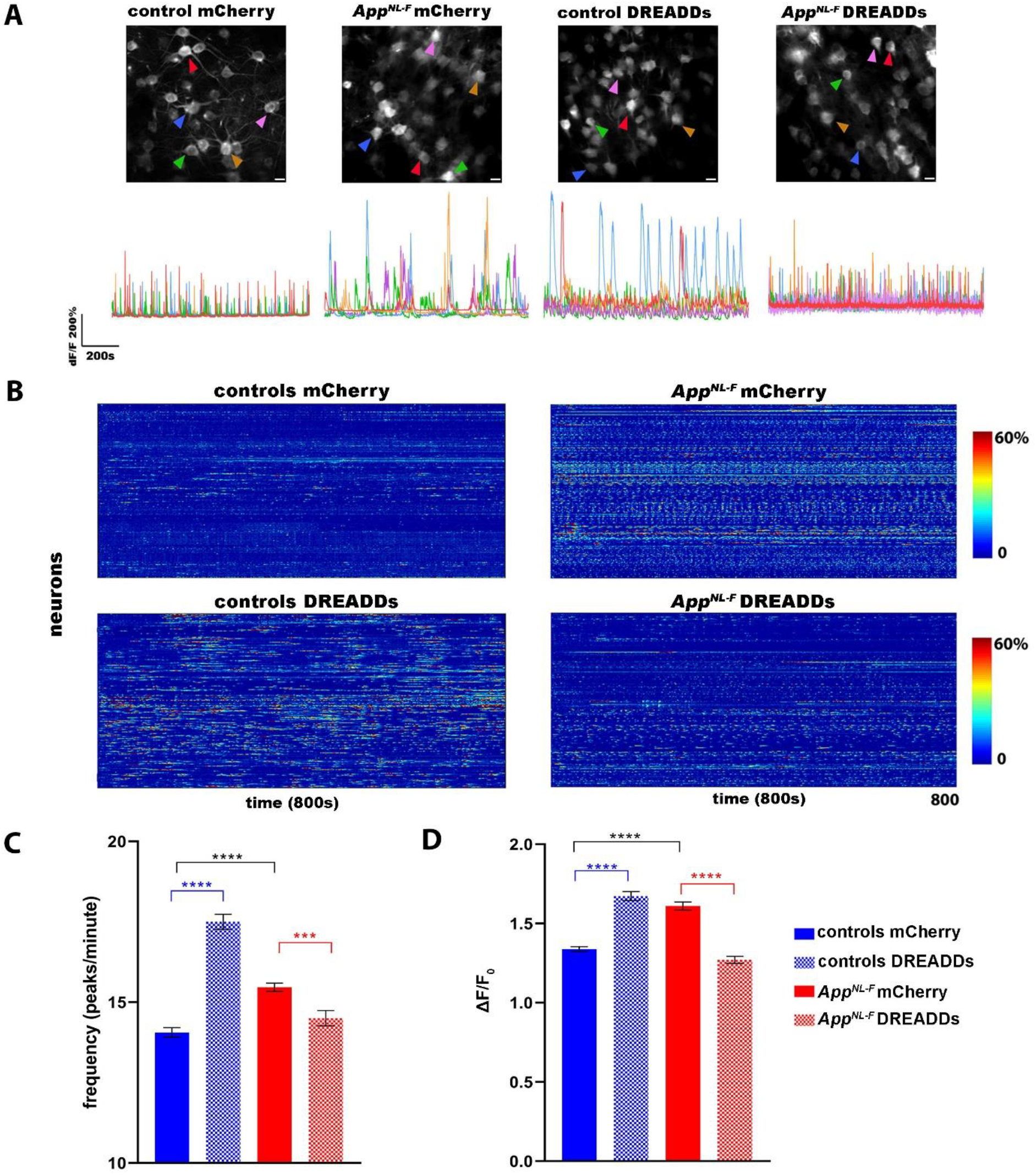
Recovery of deficient calcium signaling in astrocytes dampens neuronal hyperactivity. **A)** Representative ΔF/F0 calcium traces of neurons in control and *App^NL-F^* mice expressing neuronal GCaMP6f (hSYN promoter) in addition to either control AAV (mCherry) or DREADDs in astrocytes (GFAP promoter). Scale bar=20μm. **B)** Heat maps show calcium activity (>20% baseline, Y-axis) over time (800s, X-axis), quantifying neuronal calcium signals in control mice expressing mCherry (N=5 mice, 463 neurons), or DREADDs (N=4 mice, 640 neurons), and *App^NL-F^* mice expressing mCherry (N=5 mice, 652 neurons) or DREADDs (N=5 mice, 466 neurons). **C-D)** Graphs show frequency (number of peaks per minute ±SEM) and signal amplitude (ΔF/F0 ±SEM). ***p<0.001, ****p<0.0001, one-way ANOVA with Sidak correction.

It is striking that the activation of DREADDs in astrocytes elicits a different response on neuronal activity in control versus *App^NL-F^* mice. We hypothesized that astrocytes regulate neuronal activity and adapt their response according to the ongoing excitation state of the neuronal network. We confirm that DREADDs-mediated stimulation of astrocytes in healthy C57BL/6 mice, i.e. with normal excitation states, shifts the balance towards increased neuronal excitation. We then asked whether astrocytic DREADD activation would change towards a more inhibitory effect if we first pharmacologically induced neuronal hyperactivity using a GLT-1 inhibitor, dihydrokainic acid (DHK, 10mg/kg). We found that DREADDs activation in healthy astrocytes decreases neuronal hyperactivity induced by DHK, recovering the hyperexcitation to baseline levels (**Fig. S5).** These data demonstrate that astrocytic calcium signaling acutely regulates changes of neuronal excitation in the healthy brain to restore network homeostasis. We conclude that such homeostatic mechanism is affected in *App^NL-F^* mice and that recovering calcium signaling in these defective astrocytes enables them to regulate neuronal hyperactivity.

### Recovery of astrocyte dysfunction mitigates clinically relevant phenotypes

Our rsfMRI data confirms earlier results **(Fig. S4)** showing that activation of astrocytes in control mice increased BOLD FC (z-scores for controls mCherry 0.19 ± 0.018 and controls DREADDs 0.31 ± 0.024, p=0.0002, N=8/group). Importantly, DREADDs-mediated astrocyte activation normalizes BOLD FC in *App^NL-F^* mice (z-score for *App^NL-F^* mCherry 0.31 ± 0.022 and *App^N-LF^* DREADDs 0.18±0.0079, p=0.0001, N=8/group) **(Fig. 4A-B),** which is also consistent with our neuronal calcium imaging data **(Fig. 3).** Hence, we show that restoring local astrocyte activity in *App^NL-F^* mice is sufficient to mitigate hypersynchrony of BOLD signals in the cingulate cortex. Interestingly, we found that worsening the astrocyte deficit in 3 months old *App^NL-F^* mice by overexpressing p130PH exacerbates BOLD hypersynchrony (**Fig. S6**), providing further support to our findings.

**Fig. 4:**
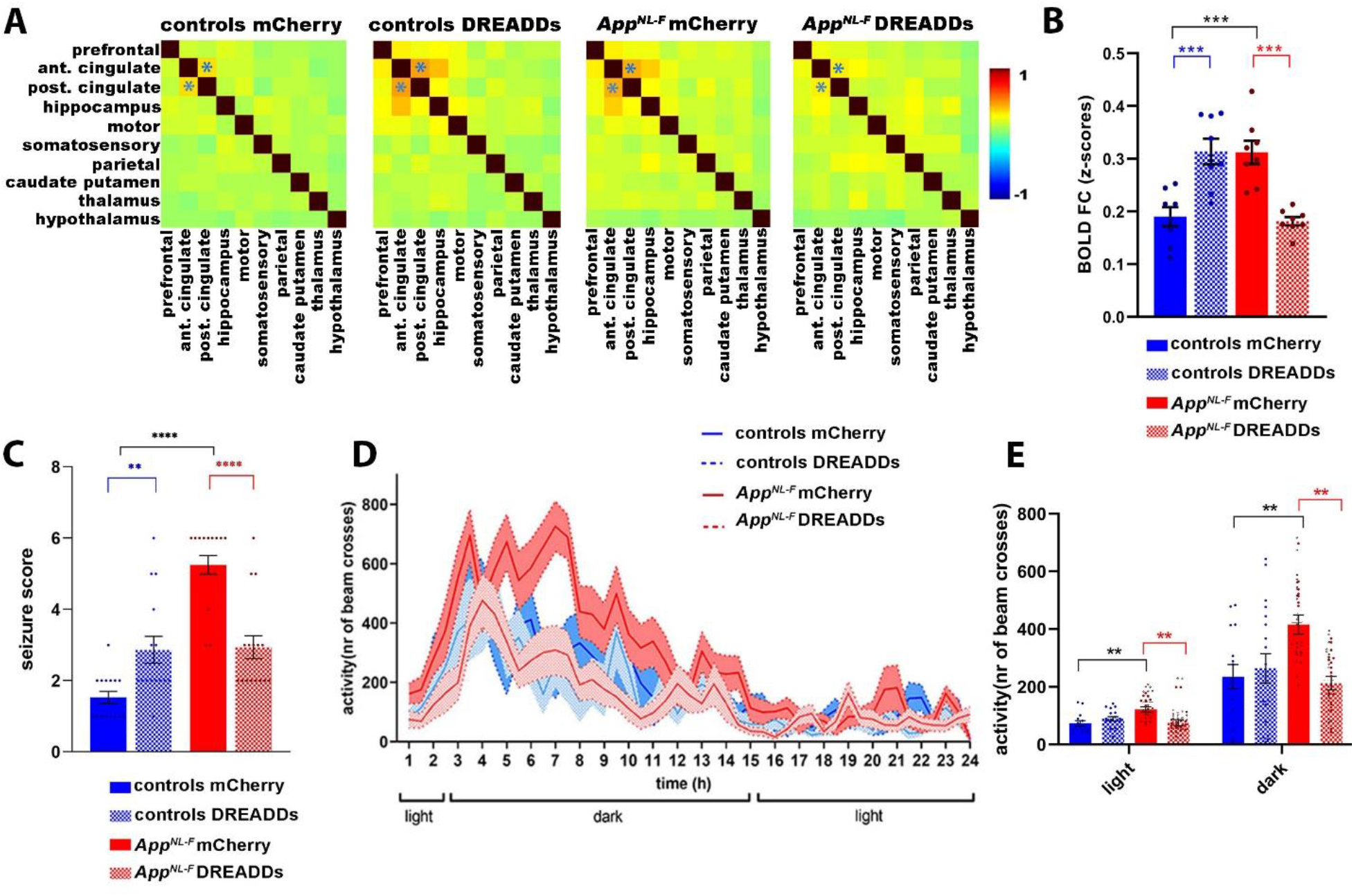
Recovery of deficient astrocyte signaling mitigates network hypersynchrony and behavior hyperactivity in early AD. **A-B)** BOLD FC matrices (**A**) and graphs quantifying FC (z-scores ±SEM) (**B**) between the anterior and posterior cingulate cortex (indicated on matrices by *) upon modulation of astrocyte calcium activity in control and *App^NL-F^* mice expressing mCherry or DREADDs (GFAP promoter) (N=8/group). ***p<0.001, ****p<0.0001, one-way ANOVA with Sidak correction. **C)** Seizure susceptibility (seizure score ± SEM) to a sub-convulsive dose of PTZ (30mg/kg) for control (N=15/group) and *App^NL-F^* mice (N=16/group) expressing mCherry or DREADDs. *p<0.05, ****p<0.0001, one-way ANOVA with Sidak correction. **D-E)** Behavior analyses of day/night activity of control and *App^NL-F^* mice expressing mCherry (control mCherry N=15, *App^NL-F^* mCherry N=16) or DREADDs (control DREADDs N=18, *App^NL-F^* DREADDs N=19) over 24 hours (counts of photobeam crosses±SEM). ****p<0.0001, one-way ANOVA with Sidak correction.

AD patients develop subclinical epileptic activity and sleep disturbances before major cognitive decline, both of which could be related to neuronal hyperactivity (23–25).

We assessed seizure susceptibility(26) in 3 months old *App^NL-F^* mice (**Fig. 4C**) by administering sub-convulsive doses of pentylene-tetrazole (27) (PTZ, 30mg/kg). Activating DREADDs in astrocytes of control mice prior to PTZ injection increases the seizure score (controls mCherry 1.5±0.16, controls DREADDs 2.9±0.37, p=0.003, N=15/group). *App^NL-F^* mice (N=16/group) have a higher seizure score compared to control mice *(App^NL-F^* mCherry 5.2±0.27, p<0.0001), which we could recover by DREADDs-mediated stimulation of astrocytes *(App^NL-F^* DREADDs 2.9±0.32, p<0.0001).

Similarly, we observed behavior hyperactivity using activity cage monitoring in 3 months old *App^NL-F^* mice (**Fig. 4D-E**). During both light and dark phases *App^NL-F^* mice show increased activity compared to controls (light: number of beam crosses for controls mCherry 73±8.8 for N=15 mice, *App^NL-F^* mCherry 121±9.37 for N=16 mice, p=0.002; dark: controls mCherry 235 ±42.1, *App^NL-F^* mCherry 416 ±33.2, p=0.005). Activation of DREADDs has no effect in control mice (light: number of beam crosses for control DREADDs 89±7.5 for N=18 mice, p=0.25; dark: controls DREADDs 264±53.1, p=0.6), but recovers behavior hyperactivity in *App^NL-F^* mice (light: number of beam crosses for *App^NL-F^* DREADDs 75±10 for N=19 mice, p=0.002; dark: *App^NL-F^* DREADDs 212±23.5, p=0.001).

Taken together, these data indicate that clinically relevant phenotypes that occur at early stages of AD, before amyloid plaque deposition, can be prevented by manipulating astrocyte calcium signaling.

## Discussion

The current study provides novel key findings on the role of astrocytes in early network hyperactivity, which is one of the earliest functional changes observed in several amyloid mouse models (2, 3) and MCI patients (28, 29). We used whole-brain resting-state functional MRI as a translational imaging tool to pinpoint brain areas with altered network FC before visible amyloid deposition in humans and mice, and identified the cingulate cortex as an early responder to AD pathology in both species. Although previous studies have reported increased network FC in patients at risk for developing AD (6), we show here that increased FC of the cingulate cortex occurs in cognitively normal individuals *preceding* the accumulation of amyloid and *predicting* amyloid-PET positivity several years later, suggesting that this is one of the earliest symptoms of incipient AD. Moreover, we show comparable increases of FC in the *App^NL-F^* mice, emphasizing the clinical relevance of this mouse model.

In-vivo studies on astrocytes in AD models are scarce, but show increased calcium signals in astrocytes of APP/PS1 mice *after* amyloid plaque deposition(14, 30). To the best of our knowledge, we are the first to report decreased *in-vivo* calcium signaling in astrocytes, several months *before* the presence of amyloid plaques. Moreover, we show decreased expression of the IP3R2 in *App^NL-F^* mice, which could explain impaired calcium release from internal stores. IP3R2 expression was also decreased in human AD brain tissue, emphasizing the translational relevance of the observations in the *App^NL-F^* mouse model. At the functional level, we confirm the importance of the IP3 signaling pathway through recovery of astrocyte calcium signaling in *App^NL-F^* mice using DREADDs.

Recovery of deficient calcium signaling in cingulate cortex astrocytes of *App^NL-F^* mice normalizes neuronal hyperactivity and increased FC. This is a major finding that emphasizes the important role of astrocytes during the earliest stages of AD. Our data demonstrate that important regulatory functions of healthy astrocytes are affected in early AD, which could maintain pathological network hyperactivity. Indeed, stimulating healthy astrocytes after pharmacologically induced neuronal hyperactivity leads to dampening of neuronal excitation. Moreover, using widefield calcium imaging and rsfMRI, we show that astrocytes maintain FC in the healthy brain.

Interestingly, recovery of astrocyte calcium signals in the anterior cingulate cortex was sufficient to rescue seizure susceptibility and behavior hyperactivity in 3 months old *App^NL-F^* mice. The anterior cingulate cortex is involved in many cognitive functions, such as learning and memory, cognitive flexibility, decision making, but also in social interactions and emotional functions through strong structural and functional connections with the posterior cingulate cortex, hippocampus, prefrontal cortex, and thalamus(31, 32). Furthermore, this region is a known epileptogenic zone (33). Our experiments highlight the importance of the cingulate cortex and the consequences of its vulnerability in early AD at the cell-, network- and behavior level (34, 35).

In conclusion, astrocytes represent a major upstream player in early AD pathology. Restoring the calcium dependent homeostatic regulatory mechanism in astrocytes mitigates early, clinically relevant phenotypes linked to network hyperactivity in AD.

## Acknowledgements

DS receives funding from the Fund for Scientific Research Flanders (FWO) (postdoctoral research fellowship, grant agreement nr. 12R1122N), FWO Krediet aan Navorsers (grant nr. 1502020N), the Alzheimer’s Association (Alzheimer’s Association Research Fellowship, grant nr. 2019-AARF-640959), and European Molecular Biology Organization (EMBO fellowship, grant nr. 7806). JW received funding from FWO as a postdoctoral grant (grant nr. 12V7519N and 12V7522N) and Krediet aan Navorsers (grant nr. 1513020N). This research received funding from the European Research Council (ERC) grant CELLPHASE_AD834682 (EU), grants from KU Leuven, VIB, Stichting Alzheimer Onderzoek Belgium (SAO), the UCB grant from the Elisabeth Foundation, a Methusalem grant from KU Leuven and the Flemish Government, and the Dementia Research Institute - MRC (UK). BDS is the Bax-Vanluffelen Chair for Alzheimer’s Disease and is supported by the Opening the Future campaign and Mission Lucidity of KUL, Leuven University. DRT receives funding from FWO (G0F8516N, G065721N). DRT received speaker honorarium from Biogen (USA), and collaborated with GE-Healthcare (UK), Novartis Pharma Basel (Switzerland), Probiodrug (Germany), and Janssen Pharmaceutical Companies (Belgium).

## Author’s contributions

Research conceptualization: DS, BDS; Methodology design: DS, BDS, UH, VB, RD, ZCV, DRT, RV, MH; Generation mouse lines: VH; Data acquisition: DS, ESL, TJ, PP, DM, TJ, KC, KH, LW; Data analysis: DS, WG, JW, MF; Technical support: WG, BV. DS and BDS wrote the manuscript with input from all authors. All authors reviewed and approved the manuscript

## Supplementary data

Supplementary methods

Supplementary figures S1-S6

Supplementary tables S1-S2

## References

1. Palop JJ, Mucke L (2010) Amyloid-B-induced neuronal dysfunction in Alzheimer’s disease: From synapses toward neural networks. Nat Neurosci 13(7):812–818.

2. Busche MA, Konnerth A (2015) Neuronal hyperactivity--A key defect in Alzheimer’s disease? Bioessays 37(6):624–32.

3. Shah D, et al. (2016) Early pathologic amyloid induces hypersynchrony of BOLD resting-state networks in transgenic mice and provides an early therapeutic window before amyloid plaque deposition. Alzheimer’s Dement 12(9):964–976.

4. Shah D, Latif-Hernandez A, De Strooper B, Saito T, Saido T, Verhoye M, D’Hooge R, Van der Linden A (2018) Spatial reversal learning defect coincides with hypersynchronous telencephalic BOLD functional connectivity in APPNL-F/NL-F knock-in mice. Sci Rep. doi:doi:10.1038/s41598-018-24657-9.

5. Hernandez AL, et al. (2017) Subtle behavioral changes and increased prefrontal-hippocampal network synchronicity in APP NL-G-F mice before prominent plaque deposition. Behav Brain Res. doi:10.1016/j.bbr.2017.11.017.

6. Kucikova L, et al. (2021) Resting-state brain connectivity in healthy young and middle-aged adults at risk of progressive Alzheimer’s disease. Neurosci Biobehav Rev 129. doi:10.1016/j.neubiorev.2021.07.024.

7. Arranz AM, De Strooper B (2019) The role of astroglia in Alzheimer’s disease: pathophysiology and clinical implications. Lancet Neurol 18(4). doi:10.1016/S1474-4422(18)30490-3.

8. Khakh BS, Deneen B (2019) The Emerging Nature of Astrocyte Diversity. Annu Rev Neurosci. doi:10.1146/annurev-neuro-070918-050443.

9. Lines J, Martin ED, Kofuji P, Aguilar J, Araque A (2020) Astrocytes modulate sensory-evoked neuronal network activity. Nat Commun 11(1). doi:10.1038/s41467-020-17536-3.

10. Poskanzer K, et al. (2018) Functional Roles of Astrocyte Calcium Elevations: From Synapses to Behavior DIFFERENT FORMS OF CALCIUM ELEVATIONS IN ASTROCYTES: THE COMPLEXITY OF SIMPLICITY. Front Cell Neurosci / www.frontiersin.org.

11. Haydon PG (2001) Glia: Listening and talking to the synapse. Nat Rev Neurosci 2(3). doi:10.1038/35058528.

12. Pérez-Alvarez A, Araque A (2013) Astrocyte-Neuron Interaction at Tripartite Synapses. Curr Drug Targets 14(11). doi:10.2174/13894501113149990203.

13. Shigetomi E, Saito K, Sano F, Koizumi SC (2019) Aberrant calcium signals in reactive astrocytes: A key process in neurological disorders. Int J Mol Sci 20(4). doi:10.3390/ijms20040996.

14. Kuchibhotla K V., Lattarulo CR, Hyman BT, Bacskai BJ (2009) Synchronous hyperactivity and intercellular calcium waves in astrocytes in Alzheimer mice. Science (80-) 323(5918):1211–1215.

15. Saito T, et al. (2014) Single App knock-in mouse models of Alzheimer’s disease. Nat Neurosci 17(5):661–663.

16. Schaeverbeke JM, et al. (2021) Baseline cognition is the best predictor of 4-year cognitive change in cognitively intact older adults. Alzheimer’s Res Ther 13(1). doi:10.1186/s13195-021-00798-4.

17. Karran E, De Strooper B (2022) The amyloid hypothesis in Alzheimer disease: new insights from new therapeutics. Nat Rev Drug Discov. doi:https://doi.org/10.1038/s41573-022-00391-w.

18. Roth BL (2016) DREADDs for Neuroscientists. Neuron 89(4):683–694.

19. Xie Y, Wang T, Sun GY, Ding S (2010) Specific disruption of astrocytic Ca2+ signaling pathway in vivo by adeno-associated viral transduction. Neuroscience 170(4):992–1003.

20. Sharp AH, et al. (1999) Differential cellular expression of isoforms of inositol 1,4,5-triphosphate receptors in neurons and glia in brain. J Comp Neurol 406(2). doi:10.1002/(SICI)1096-9861(19990405)406:2<207::AID-CNE6>3.0.CO;2-7.

21. Taylor CW, Genazzani AA, Morris SA (1999) Expression of inositol trisphosphate receptors. Cell Calcium 26(6). doi:10.1054/ceca.1999.0090.

22. Busche MA, et al. (2008) Clusters of hyperactive neurons near amyloid plaques in a mouse model of Alzheimer’s disease. Science (80-) 321(5896):1686–1689.

23. Vossel KA, et al. (2013) Seizures and epileptiform activity in the early stages of Alzheimer disease. JAMA Neurol 70(9). doi:10.1001/jamaneurol.2013.136.

24. Lam AD, et al. (2017) Silent hippocampal seizures and spikes identified by foramen ovale electrodes in Alzheimer’s disease. Nat Med 23(6). doi:10.1038/nm.4330.

25. Ju YES, Lucey BP, Holtzman DM (2014) Sleep and Alzheimer disease pathology-a bidirectional relationship. Nat Rev Neurol 10(2). doi:10.1038/nrneurol.2013.269.

26. Van Erum J, Van Dam D, De Deyn PP (2019) PTZ-induced seizures in mice require a revised Racine scale. Epilepsy Behav 95. doi:10.1016/j.yebeh.2019.02.029.

27. Shimada T, Yamagata K (2018) Pentylenetetrazole-induced kindling mouse model. J Vis Exp 2018(136). doi:10.3791/56573.

28. Ewers M, Sperling RA, Klunk WE, Weiner MW, Hampel H (2011) Neuroimaging markers for the prediction and early diagnosis of Alzheimer’s disease dementia. Trends Neurosci 34(8). doi:10.1016/j.tins.2011.05.005.

29. Bakker A, et al. (2012) Reduction of Hippocampal Hyperactivity Improves Cognition in Amnestic Mild Cognitive Impairment. Neuron 74(3). doi:10.1016/j.neuron.2012.03.023.

30. Reichenbach N, et al. (2018) P2Y1 receptor blockade normalizes network dysfunction and cognition in an Alzheimer’s disease model. J Exp Med 215(6). doi:10.1084/jem.20171487.

31. Stevens FL, Hurley RA, Taber KH (2011) Anterior Cingulate Cortex: Unique Role in Cognition and Emotion. J Neuropsychiatry Clin Neurosci 23(2):121–125.

32. Allman JM, Hakeem a, Erwin JM, Nimchinsky E, Hof P (2001) The anterior cingulate cortex. The evolution of an interface between emotion and cognition. Ann N Y AcadSci 935:107–117.

33. Chang WP, Shyu BC (2014) Anterior cingulate epilepsy: Mechanisms and modulation. Front Integr Neurosci 7(JAN). doi:10.3389/fnint.2013.00104.

34. Borges CR, Poyares D, Piovezan R, Nitrini R, Brucki S (2019) Alzheimer’s disease and sleep disturbances: a review. Arq Neuropsiquiatr 77(11). doi:10.1590/0004-282X20190149.

35. Keszycki RM, Fisher DW, Dong H (2019) The hyperactivity-impulsivity-irritiability-disinhibition-aggression-agitation domain in Alzheimer’s disease: Current management and future directions. Front Pharmacol 10(SEP). doi:10.3389/fphar.2019.01109.

