## Supplementary methods and figures for "Astrocyte calcium dysfunction causes early network hyperactivity in Alzheimer’s Disease"

### Mice

All mice were bred in-house and maintained on a C57BL/6J background. All *AppP*$^{NL/NL}$ (KM670/671N Swedish mutation), i.e. control mice, and *App*$^{NL-F/NL-F}$ (KM670/671N Swedish and I716F Iberian mutations) knock-in mice(1) were backcrossed for at least 2 generations with C57BL/6J mice at De Strooper lab. All experimental procedures were performed in strict accordance with the European Directive 2010/63/EU on the protection of animals used for scientific purposes. The protocols were approved by the Committee on Animal Care and Use at VIB-KU Leuven, Belgium (permit number P043/2018 and P072/2019), and all efforts were made to minimize animal suffering.

### Viral vectors injections

Designer Receptors Exclusively Activated by Designer Drugs (DREADDs) were used to increase calcium signaling in astrocytes(2, 3). Overexpression of the Pleckstrin Homology (PH) domain of Phospholipase C (PLC)-like protein p130 (p130PH)(4), which buffers IP3 and blocks calcium release from internal stores, was used to decrease calcium signals in astrocytes. Expression occurred through intracerebral injections of adeno-associated viruses (AAVs). All AAVs were constructed and packaged by Vectorbuilder (www.vectorbuilder.com). We used the following AAVs (AAV8 serotype, >$10^{12}$ GC/ml): pAAV-GFAP-mCherry, pAAV-GFAP-hMD3(Gq)-mCherry, pAAV-GFAP-P130PH-mCherry, pAAV-GFAP-GCaMP6f, pAAV-hSYN-GCaMP6f, pAAV-hSYN-GCaMP6f(PHP-eB), pAAV-hSYN-eGFP. We allowed 3 weeks between AAV injections and readouts.

For stereotactic injections, mice were injected with buprenorphine (0.05 mg/kg), anesthetized with 2.5% isoflurane and placed in a stereotactic frame. Xylocaine (subcutaneous, 100µl, 1%) was used as local anesthesia before exposing the skull. A surgical drill was used to perform a small craniotomy to allow microinjection of AAVs. Bilateral injections were carried out using a 2µl neurosyringe with a 30-gauge needle (https://www.hamiltoncompany.com/) at the following coordinates: anteroposterior (AP) -1.1mm from Bregma; mediolateral (ML) ±0.3mm; dorsoventral (DV) -1.3mm, to target the anterior cingulate cortex (**Fig. S1)**. An injection pump controlled the injection volume (2X10^13 GC/ml, 0.4 µl per AAV per injection site) and rate (0.1µl/minute). After each injection, the syringe was left in place for 10 minutes to allow for diffusion of the AAVs, after which it was carefully withdrawn. The burr hole was filled with bone wax, after which the incision was closed using surgical glue (Dermafuse). All AAVs were injected stereotactically in the cingulate cortex, except for PHP.eB, which was injected into the tail vein to express

GCaMP6f in neurons. Postoperative care included subcutaneous injections of buprenorphine (0.05mg/kg) at 4-6 hours after surgery.

**Fig. S1** shows expression of AAVs in astrocytes, i.e. co-staining with GFAP (astrocytes) but not with Iba1 (microglia) or NeuN (neurons). To ascertain that AAVs targeted to modulate astrocytes are not expressed in neurons we co-injected pAAV-GFAP-mCherry (targeted to astrocytes) and pAAV-SYN-eGFP (targeted to neurons) and found no correlation between the expressions of these viruses (**Fig.S1**).

**Cranial window surgeries for multi-photon imaging**

Craniotomy surgeries were performed to gain optical access to the cingulate cortex through a set of cover glasses (5). All mice were anaesthetized using a mix of ketamine and xylazine (100 mg/kg and 10 mg/kg respectively, intraperitoneal) and placed in a stereotactic frame. Xylocaine (subcutaneous, 100µl, 1%) was used as local anesthesia before exposing the skull. A custom-made titanium head plate was mounted to the skull using adhesive cement C&B-metabond, Parkell) and acrylic material (TAB 2000, Kerr). A 6mm diameter craniotomy was made over the brain midline using a surgical drill. Cranial windows were constructed by bonding two 6mm cover glass slips to an 8 mm top cover glass using optical adhesive (NOA 71, Norland), implanted over the craniotomy and sealed using Vetbond (3M). The top layer was covered with acrylic material mixed with black pigment to block ambient light from the two-photon microscope. Two rubber rings were attached to the head-plate to form a well to hold distilled water during imaging. Buprenorphine was administered postoperatively (0.05 mg/kg) when the animal recovered from anesthesia every 12 hours until 3 days after surgery.

**Two-photon calcium imaging**

During all imaging procedures mice were anesthetized using a combination of medetomidine (Domitor, 0.05mg/kg, subcutaneous) and isoflurane (0.5%), according to a previously established protocol(6–8). After the imaging procedures, the effects of medetomidine were counteracted by atipamezole (Antisedan, 0.1mg/kg).

A customized two-photon microscope (Neurolabware) was used for all imaging. GCaMP6f was excited at 920 nm wavelength with a Ti:Sapphire excitation laser (MaiTai DeepSee, Spectra-Physics). The emitted photons were split by a dichroic beamsplitter (centered at 562 nm) and collected with a photomultiplier tube (PMT, Hamamatsu) through a bandpass filter (510 ± 42 nm, Semrock) for the green fluorescence of GCaMP and a bandpass filter (607 ± 35 nm, Semrock) for the red fluorescence of mCherry. Two-photon images (512x512 pixels per frame) were collected at 30 Hz with a 16x objective (Nikon 0.80 NA) for 12-15 minutes to measure spontaneous calcium activity in astrocytes or neurons (imaging depth 150-300μm). Clozapine-N-oxide (CNO, 3mg/kg, intraperitoneal) was used to activate DREADDs by a single injection 20 minutes before imaging. For the experiments with dihydrokainic acid (DHK), we acquired 5 minutes long scans at baseline, after DHK (10mg/kg, intraperitoneal) or saline injections, and after subsequent CNO (3mg/kg) injections.

All data were analyzed using Image J/Fiji software (https://imagej.nih.gov) and custom written scripts in MATLAB 2019b (Mathworks, Natick, MA, USA) as previously established(9). Two-photon images for all experiments were motion registered, and regions-of-interest (ROIs) were manually segmented. We analyzed somas and proximal processes for astrocytes and somas for neurons. Cellular time courses for each ROI were extracted by averaging all pixels inside each ROI and removing the background. All time traces were then converted to $\Delta F/F_0$ traces. For quantification, we defined events as $\Delta F/F_0$ signals that exceeded twice the standard deviation of the noise band. We then quantified the number of transients and $\Delta F/F_0$ amplitude for each group (9). Statistical analyses included one-way ANOVAs with Sidak correction for multiple comparisons to compare multiple groups.

**Widefield imaging**

Widefield fluorescent images were acquired through a 2x objective (NA = 0.055, Edmund Optics) using blue LED (479 nm, ThorLabs) and collection of the emitted green light with an EMCCD camera (510/84 nm filter, Semrock) at a rate of 20 frames/second. CNO (3mg/kg) was injected in all experimental groups 20 minutes before imaging. For the analysis, images were rescaled to 2/3 of their original size, co-registered, and converted to $\Delta F/F_0$ using Image J according to an established protocol(8). ROIs were defined along the midline and lateral parts of the window and were then used for correlation analyses, where pairwise Pearson correlation coefficients between each pair of ROIs were calculated and z-transformed using MATLAB. Mean z-transformed FC matrices were calculated for each group. Additionally, seed-based analyses were performed by computing individual z-transformed FC-maps of the left cingulate cortex, resulting in zFC-maps for each group. Statistical analyses included one or two-way ANOVAs comparing functional connectivity (z-score) between the anterior and posterior cingulate cortex, or between lateral cortical areas, with Sidak correction for multiple comparisons.

**Mouse resting state functional Magnetic Resonance Imaging**

During all imaging procedures mice were anesthetized using a combination of medetomidine (Domitor, 0.05mg/kg, subcutaneous) and isoflurane (0.5%), according to a previously established protocol(6, 7).

After the imaging procedures, the effects of medetomidine were counteracted by atipamezole (Antisedan, 0.1mg/kg).

MRI procedures were performed on a 9.4T Biospec MRI system (Bruker BioSpin, Germany) with the Paravision 6 software (www.bruker.com). Images were acquired using a standard Bruker cross coil set-up with a quadrature volume transmit coil and a quadrature surface receive coil for mice. Three orthogonal multi-slice Turbo RARE T2-weighted images were acquired to render slice-positioning uniform (repetition time 2000 ms, echo time 33 ms, 16 slices of 0.4 mm with a gap of 0.1 mm). Field maps were acquired for each animal to assess field homogeneity, followed by local shimming, which corrects for the measured inhomogeneity in a rectangular VOI within the brain. Resting-state signals were measured using a T2*-weighted single shot echo-planar imaging sequence (repetition time 2000 ms, echo time 15 ms, 16 slices of 0.4 mm with a gap of 0.1 mm, 300 repetitions). The field-of-view was (20 x 20) $mm^2$ and the matrix size (128 x 64), resulting in voxel dimensions of (0.156 x 0.312 x 0.5) $mm^3$. CNO (1, 3, or 10 mg/kg, intraperitoneal) was injected 20 minutes before rsfMRI imaging sessions.

Pre-processing of the rsfMRI data, including realignment, normalization and smoothing, was performed using SPM12 software (Statistical Parametric Mapping, http://www.fil.ion.ucl.ac.uk) as described previousl (6). First, all images within each session were realigned to the first image. For the rsfMRI data analyses, motion parameters resulting from the realignment were included as covariates to correct for variation in intensity related to possible movement that occurred during the scanning procedure. Second, all datasets were normalized to a study specific EPI template. The EPI images were co-registered to a study-specific anatomical T2-weighted template and the Allen mouse brain atlas (https://mouse.brain-map.org/static/atlas). Finally, in plane smoothing was done using a Gaussian kernel with full width at half maximum of twice the voxel size (0.31 X 0.62 $mm^2$). All rsfMRI data were filtered between 0.008-0.1 Hz.

ROIs were defined using MRicron software (https://www.nitrc.org/projects/mricron) and the Allen Brain Atlas. ROI-correlation analyses were performed using the MATLAB REST toolbox (http://resting-fmri.sourceforge.net), resulting in z-transformed FC matrices. Additionally, seed-based analyses were performed by computing individual z-transformed FC-maps of the left cingulate cortex, resulting in mean zFC-maps for each group (FDR-corrected, $p<0.05$, threshold of 20 voxels). Statistical analyses included two-sample T-tests to compare two groups. For the comparison of FC matrices and FC maps, we used FDR correction for multiple comparisons ($p<0.05$). One-way ANOVA with Sidak correction was used to compare multiple groups.

**Human Positron Emission Tomography and resting state functional Magnetic Resonance Imaging**

***F-PACK participants***

The Flemish Prevent Alzheimer's Disease Cohort KU Leuven (F-PACK) is a cohort of 180 participants followed by the Laboratory for Cognitive Neurology, KU Leuven. Individuals were recruited between 2009 and 2015 in three waves of 60. Inclusion criteria included participants being aged between 50-80 years old, scoring ≥ 27 on the mini mental state examination, having a clinical dementia rating score of 0 and scoring within published norms on an extensive neuropsychological test battery(10, 11). Individuals were excluded if they had any contraindications for magnetic resonance imaging (MRI), lesions on MRI, history of cancer or neuropsychological illness, or exposure to radiation one year preceding the baseline amyloid-PET scan. Recruitment was stratified for *APOE* status (*ε4* allele present or absent) and *BDNF* status (*66 met* allele present or absent), such that per five-year age bin each factorial cell was matched for age, sex, and education. At baseline all individuals received a structural MRI scan and an $^{18}$F-Flutemetamol amyloid-PET scan. A subset also received baseline functional MRI and follow-up amyloid-PET and thus form the cohort for the present study. Individuals are being followed over a 10-year period with two-yearly neuropsychological assessments. **Supplementary Table 1** shows details on each participant included for the analyses.

***Amyloid-PET***

Baseline and follow-up amyloid-PET scans were acquired on a 16-slice Biograph PET/CT scanner (Siemens, Erlangen, Germany). All baseline and follow-up scans were acquired as a static $^{18}$F-Flutemetamol PET scan with acquisition window 90-120 minutes post injection, with an average intravenous dose of 150 ± 1.02 MBq (139 – 164MBq) at baseline and 173 ± 3.83 MBq (140 MBq – 197 MBq) at follow-up as described previously(11–13). A subset of 14 participants received a dynamic $^{11}$C-Pittsburg Compound-B ($^{11}$C-PiB) scan lasting 60 minutes. A low-dose CT scan was acquired prior to the PET for attenuation correction, and random and scatter corrections were applied. Data were reconstructed using ordered subsets expectation maximization, as six frames of five minutes (five iterations, eight subsets). PET images were processed using SPM12 and MATLAB as described previously (11–13).

Mean standardized uptake value ratios (SUVRs) were calculated at baseline and follow-up in a composite cortical volume of interest (VOI) derived from the Automated Anatomic Labelling Atlas (AAL), with cerebellar gray matter as reference region. Frontal (AAL areas 3-10, 13-16, 23-28), parietal (AAL 57-70), anterior cingulate (AAL 31-32), posterior cingulate (AAL 35-36), and lateral temporal (AAL 81-82, 85-90) regions were included(11, 14). This VOI was masked with the participant-specific grey matter (GM) segmentation map (threshold of grey matter voxel >0.3). A subject-specific cerebellar grey matter reference region was used for the calculation of the SUVRs. This was defined as AAL 91-108 and was masked by the participant-specific GM map (GM threshold >0.3). SUVRs were then converted to Centiloids(15) using the formulas $CL_{Flut}$ = 127.6 x SUVR -149, $CL_{PiB}$ = 132.53 x SUVR – 174.64(16, 17). Individuals were included in the analysis with following criteria: baseline CL<23.5 for all individuals, CL<23.5 at follow-up (T1) for the control group, CL≥23.5 at T1 for amyloid accumulators. A CL value of 23.5 was chosen as threshold as it indicates amyloid positivity(10).

***MRI***

A 3T Philips Achieva MRI scanner with a 32-channel head coil (Philips, Best, The Netherlands) was used to obtain high resolution T1-weighted structural MRI scans at baseline and follow-up (inversion time = 900ms, repetition time = 9.6ms, echo time = 4.6ms, flip angle = 8°, field of view = 250 x 250 mm, voxel size = 0.98 x 0.98 x 1.2 $mm^3$). Eyes-closed resting-state fMRI data was acquired using T2* echo-planar images (repetition time 3000ms, echo time 30ms, 50 transverse slices, voxel size 2.5x2.5x2.5 $mm^3$, field-of-view 200x200$mm^2$)(11).

Preprocessing of rsfMRI data was performed in SPM12 as described previously(11, 18) and included realignment, co-registration of structural and functional images, normalization of structural images to the SPM12 T1-template in Montreal Neurological Institute (MNI) space, normalization of the rsfMRI scans and smoothing using a Gaussian kernel with full width at half maximum (6x6x6)$mm^3$. All data were filtered using a band pass filter of 0.008-1Hz. For the rsfMRI data analyses, the effects of motion, global signal and white matter signal were regressed out. The global signal regressor was calculated by averaging the BOLD time series across all voxels in which the sum of the grey matter/CSF/white matter segmentation maps were above 0.9. Motion parameters were derived from the realignment step.

ROIs were defined on the MNI template using MRicron software and the Brainnetome atlas (https://atlas.brainnetome.org/). ROI-correlation analyses were performed in MATLAB, resulting in z-transformed FC matrices. Regression analysis was performed to assess group differences in FC using amyloid load as a covariate. Statistical analyses included two-sample T-tests to compare two groups. For the comparison of FC matrices between two groups, we used FDR-correction for multiple comparisons

($p<0.05$). Seed-based analyses were performed by computing individual z-transformed FC-maps of the left cingulate cortex, resulting in mean zFC-maps for each group (FDR-corrected, $p<0.05$, threshold of 20 voxels). Statistical analyses included two-sample T-tests to compare FC maps between two groups (uncorrected, $p<0.001$, threshold of 20 voxels).

**Behavior studies**

For the activity measurements, CNO was administered for 24h through drinking water. All mice were housed individually in $20 \times 30$ $cm^2$ transparent cages located between three photo beams. Activity was measured as number of beam crossings with a 30 minutes interval over 24 hours. Analyses included one-way ANOVAs with Sidak correction for multiple comparisons to compare activity during light and dark phases for each group.

For the seizure susceptibility experiments, mice were injected with CNO (3mg/kg, intraperitoneal). After 30 minutes, a subconvulsive dose of pentylene-tetrazole (PTZ, 30mg/kg, intraperioneal) was administered and mice were placed individually in a recording chamber. Recording started immediately after PTZ injection and lasted 30 minutes. Seizures were scored as listed in **Supplementary Table 2** (19). Analyses included a one-way ANOVA with Sidak correction for multiple comparisons.

**Immunohistochemistry**

Mice were euthanized using intraperitoneal injection of 60 mg/kg pentobarbital, after which they were transcardially perfused first with ice-cold PBS, and then with 4% paraformaldehyde. Brains were then surgically removed, postfixed in 4% paraformaldehyde, after which 30µm thick vibratome sections were prepared. Brain sections were stored in cryoprotectant solution (30% ethylene glycol, 30% glycerol, 40% PBS) at −20 °C until use. For human samples, 10µm thick cryo-sections were made from frozen cortical tissue of AD and age-matched control samples obtained from the Netherlands Brain Bank. Immunofluorescence analyses were performed using the following antibodies: Guinea pig anti-GFAP (Synaptic Systems, ID 173004,1:1000), mouse anti-NeuN (Synaptic Systems, ID 266004, 1:200), Rabbit anti-Iba1 (Synaptic Systems, ID 234004, 1:200), Chicken anti-GFP (Abcam, ID ab13970, 1:500), Rabbit anti-RFP (Rockland, ID 600-401-379, 1:500), rabbit anti-IP3R2 (Thermofischer, ID PA1904, 1:200). Images were acquired on a NikonTi-E inverted microscope equipped with an A1R confocal unit driven by NIS (4.30) software. All images were acquired at 10x or 20x magnification at 2X zoom and analysed using Image J/FIJI software. For quantification of IP3R2 staining the density of staining per $100µ^2$ was calculated from maximum intensity projection images.

**Supplementary figures and tables**

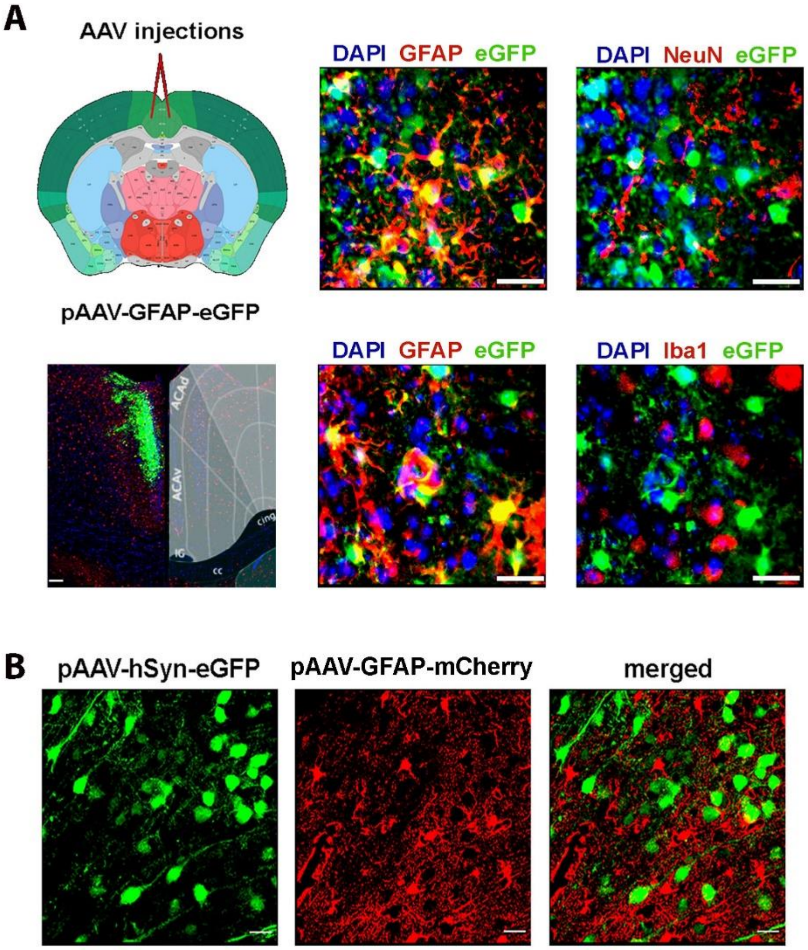


**Fig. S1: Immunohistochemistry validation of AAV injections. A)** Injection site of local AAV injections in the cingulate cortex shown on the Allen Brain Atlas. pAAV-GFAP-eGFP expression was observed in the target region, i.e. anterior cingulate cortex and there was co-staining with astrocyte marker GFAP, but not with microglia (Iba1) or neurons (NeuN). **B)** Stereotactical co-injection of pAAV-hSYN-eGFP and pAAV-GFAP-mCherry shows no co-staining, confirming specificity of each AAV.

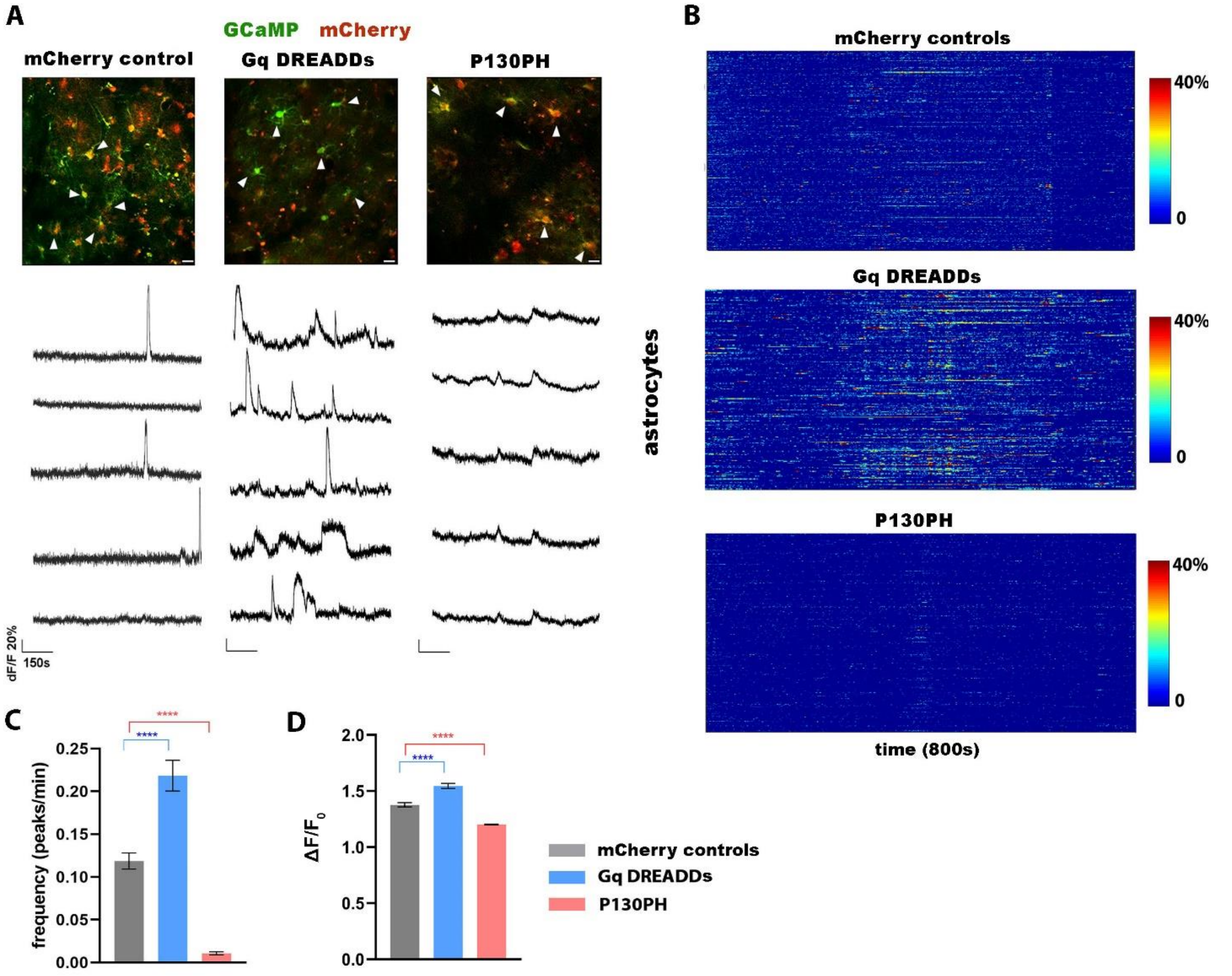


**Fig. S2: Modulation of astrocyte calcium signaling in-vivo.** Astrocytes express Gcamp6f in addition to either mCherry, DREADDs or P130PH through local AAV injections in the cingulate cortex (GFAP promoter). **A)** Representative $\Delta F/F_0$ calcium traces of astrocytes expressing mCherry, DREADDs or P130PH. DREADDs elicit increased calcium signaling in astrocytes upon CNO injection (3mg/kg) and P130PH expression causes decreased calcium signaling (4 weeks expression) compared to C57BL/6 mice expressing mCherry. Scale bar=20µm. **B)** Quantification of astrocyte calcium signals in C57BL/6 mice expressing mCherry (N=5 mice, 329 cells), DREADDs (N=5 mice, 263 cells), and P130PH (N=5 mice, 353 cells). Heat maps show calcium activity (>20% baseline, Y-axis) over time (800s, X-axis). **C-D)** Graphs show frequency (number of peaks per minute ±SEM) and signal amplitude ($\Delta F/F_0$ ±SEM), ****$p<0.0001$, one-way ANOVA with Sidak correction.

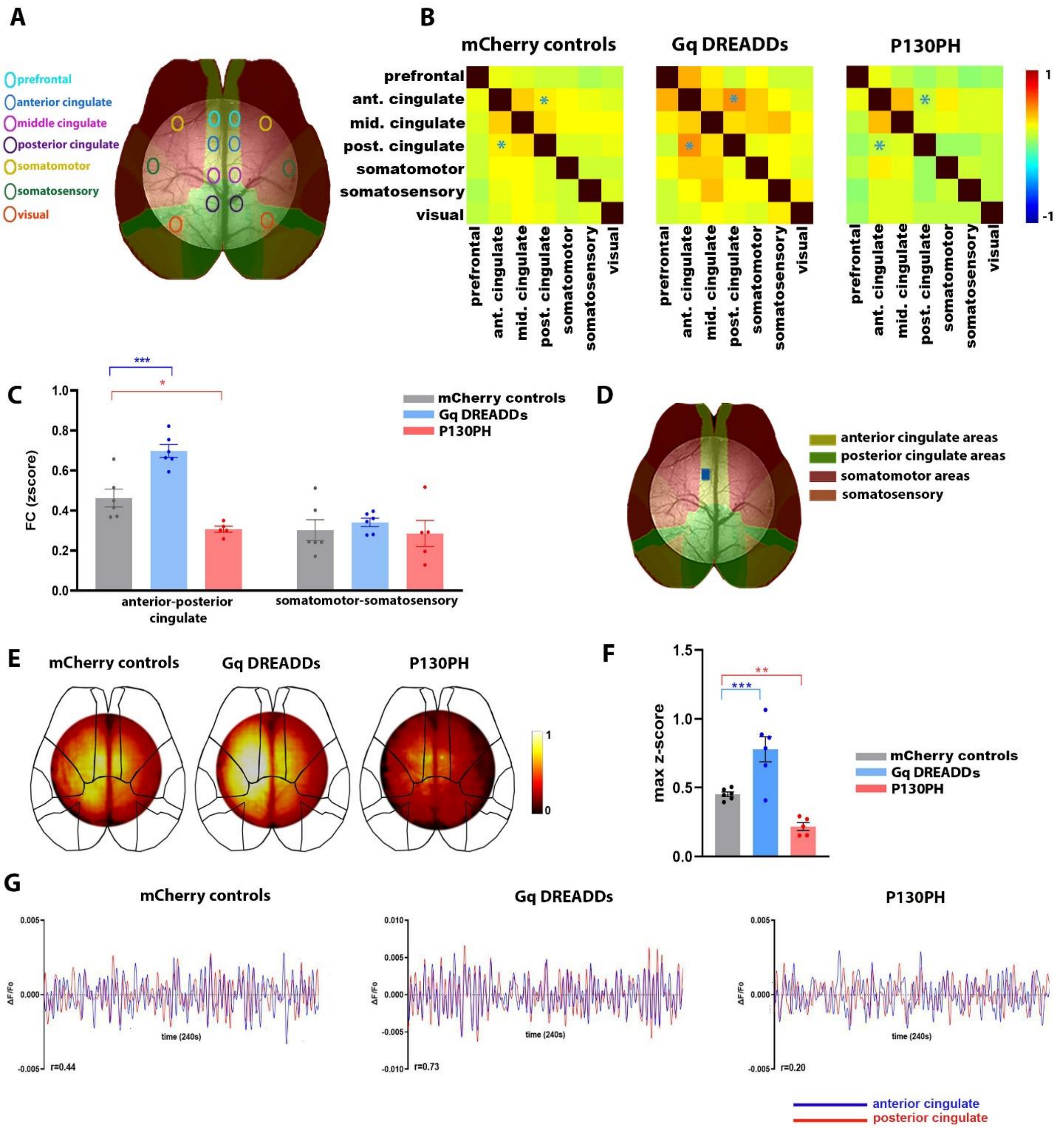
A
prefrontal
anterior cingulate
middle cingulate
posterior cingulate
somatomotor
somatosensory
visual
B
mCherry controls
Gq DREADDs
P130PH
prefrontal
ant. cingulate
mid. cingulate
post. cingulate
somatomotor
somatosensory
visual
1
-1
C
FC (zscore)
mCherry controls
Gq DREADDs
P130PH
anterior-posterior cingulate
somatomotor-somatosensory
D
anterior cingulate areas
posterior cingulate areas
somatomotor areas
somatosensory
E
mCherry controls
Gq DREADDs
P130PH
F
max z-score
mCherry controls
Gq DREADDs
P130PH
G
mCherry controls
Gq DREADDs
P130PH
ΔF/F0
time (240s)
r=0.44
r=0.73
r=0.20
anterior cingulate
posterior cingulate

**<u>Fig. S3:</u> Modulation of astrocyte calcium activity alters mesoscale neuronal network FC in C57BL/6 mice.** Astrocytes express either mCherry, DREADDs or P130PH through local AAV injections in the cingulate cortex (GFAP promoter). All mice were injected intravenously with AAV-PHP.eB to express Gcamp6f in neurons (hSYN promoter). **A)** Brain regions used for correlation analyses are shown on the Allen Mouse Brain Atlas. **B)** Mean FC matrices show correlation between $\Delta F/F_0$ calcium signals for each pair of brain regions. Color scale shows z-scores, representing strength of FC. **C)** FC between the anterior and posterior cingulate cortex (indicated by *) is increased in C57BL/6 mice expressing DREADDs (3mg/kg CNO, N=6) and decreased in mice expressing P130PH (4 weeks expression, N=5) compared to mice expressing mCherry (N=6). Graph shows FC between the anterior-posterior cingulate cortex and somatosensory-motor brain regions (z-score ± SEM). *$p<0.05$, ***$p<0.001$, two-way ANOVA with Sidak correction. **D)** Seed region (anterior cingulate cortex) used to compute FC maps shown on the mouse brain atlas. **E)** FC maps of the anterior cingulate cortex demonstrate increased cingulate FC of mice expressing DREADDs and decreased FC in mice expressing P130PH compared to mCherry control mice. Color scale shows z-scores. **F)** Graph displays maximum FC of each cingulate FC map per condition (z-max ± SEM). **$p<0.01$, ***$p<0.001$, one-way ANOVA with Sidak correction. **G)** Representative $\Delta F/F_0$ calcium time courses (time=240s) of the anterior and posterior cingulate cortex for each condition with corresponding correlation values.

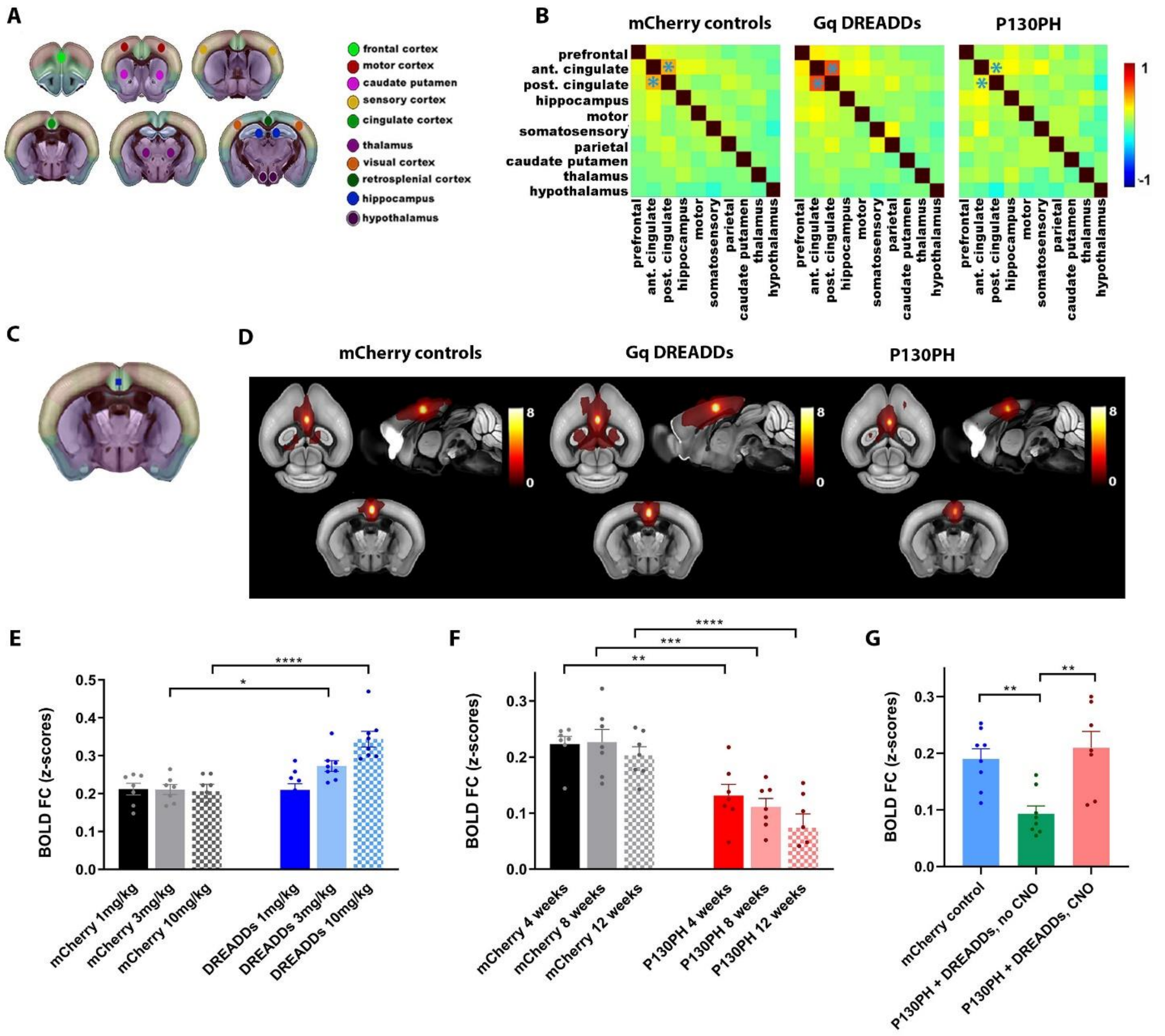

A
frontal cortex
motor cortex
caudate putamen
sensory cortex
cingulate cortex
thalamus
visual cortex
retrosplenial cortex
hippocampus
hypothalamus
B
mCherry controls
Gq DREADDs
P130PH
prefrontal
ant. cingulate
post. cingulate
hippocampus
motor
somatosensory
parietal
caudate putamen
thalamus
hypothalamus
1
-1
C
D
mCherry controls
Gq DREADDs
P130PH
8
0
E
BOLD FC (z-scores)
mCherry 1mg/kg
mCherry 3mg/kg
mCherry 10mg/kg
DREADDs 1mg/kg
DREADDs 3mg/kg
DREADDs 10mg/kg
F
BOLD FC (z-scores)
mCherry 4 weeks
mCherry 8 weeks
mCherry 12 weeks
P130PH 4 weeks
P130PH 8 weeks
P130PH 12 weeks
G
BOLD FC (z-scores)
mCherry control
P130PH + DREADDs, no CNO
P130PH + DREADDs, CNO

**Fig. S4:** **Modulation of astrocyte calcium activity alters BOLD networks in C57BL/6mice.** Astrocytes express either mCherry, DREADDs or P130PH through local AAV injections in the cingulate cortex (GFAP promoter). **A)** Brain regions used to calculate FC matrices are displayed on the Allen Brain Atlas. **B)** Mean FC matrices show correlation between BOLD signals. FC between the anterior and posterior cingulate cortex (indicated by *) is increased or decreased in C57BL/6 mice expressing DREADDs (3mg/kg CNO, N=8) or P130PH (4 weeks expression, N=7), respectively, compared to mice expressing mCherry (N=10). Color scale represents BOLD FC strength. **C)** The location of the seed region (anterior cingulate cortex) used to compute FC maps is shown on the mouse atlas. **D)** Mean BOLD FC maps of the cingulate cortex of mice expressing mCherry (N=10), DREADDs (3mg/kg CNO, N=8) or P130PH (4 weeks expression, N=7). Color scale shows T-values, which represent strength of BOLD FC. **E-F-G)** Quantification of BOLD FC between the anterior and posterior cingulate cortex (z-score ± SEM) shows a dose-dependent response of CNO (1,3 and 10 mg/kg, N=7-9/group) (**E**) and lasting effect of P130PH (4, 8- and 12-weeks expression, N=7/group) (**F**). Mice co-injected with DREADDs and P130PH (N=7/group) show decreased FC (z-score ± SEM) before CNO injection, which can be recovered upon CNO administration (3mg/kg) (**G**). *$p<0.05$, **$p<0.01$, ***$p<0.001$, ****$p<0.0001$, one-way ANOVA with Sidak correction.

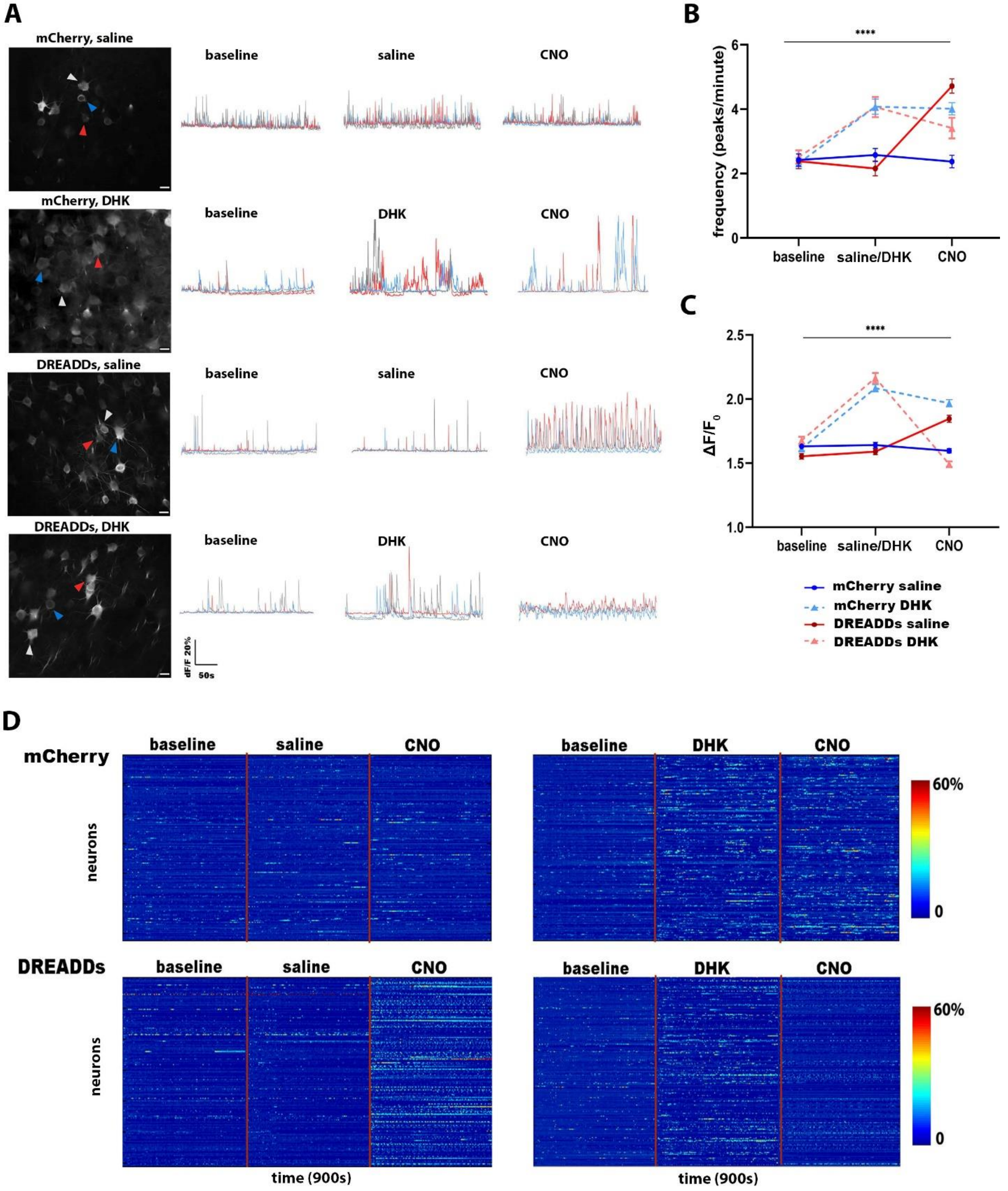
A
mCherry, saline
baseline
saline
CNO
mCherry, DHK
baseline
DHK
CNO
DREADDs, saline
baseline
saline
CNO
DREADDs, DHK
baseline
DHK
CNO
dF/F 20%
50s
B
****
frequency (peaks/minute)
6
4
2
0
baseline
saline/DHK
CNO
C
****
ΔF/F0
2.5
2.0
1.5
1.0
baseline
saline/DHK
CNO
mCherry saline
mCherry DHK
DREADDs saline
DREADDs DHK
D
mCherry
baseline
saline
CNO
baseline
DHK
CNO
neurons
60%
0
DREADDs
baseline
saline
CNO
baseline
DHK
CNO
neurons
60%
0
time (900s)
time (900s)

**Fig. S5: Astrocytes regulate neuronal activity in the healthy brain.** Astrocytes express either mCherry or DREADDs (GFAP promoter) and neurons express GCaMP6f (hSYN promoter) through local AAV injections in the cingulate cortex. **A)** Representative images and $\Delta F/F_0$ calcium time traces of neurons from C57BL/6 mice that were injected with dihydrokainic acid (DHK, 10mg/kg) or saline prior to injection of CNO (3mg/kg) to activate DREADDs in astrocytes (N=4/group, mCherry saline N=302 neurons, mCherry DHK 290 neurons, DREADDs saline N=384 neurons, DREADDs DHK N=200 neurons). Scale bar=20µm. **B-C)** Frequency (number of peaks/min ±SEM) and amplitude ($\Delta F/F_0$ ±SEM) for each condition. ****$p<0.0001$ group effect, ****$p<0.0001$, time effect, two-way ANOVA with Sidak correction. No significant differences were found for $\Delta F/F_0$ amplitude or frequency in mCherry mice injected with saline. Significant differences were observed in mCherry mice injected with DHK ($\Delta F/F_0$ and frequency: baseline versus DHK $p<0.0001$), while subsequent injection of CNO did not cause changes. Significant differences were observed for mice expressing DREADDs injected with CNO ($\Delta F/F_0$ and frequency, baseline or saline versus CNO, $p<0.0001$). Finally, significant differences were found for mice expressing DREADDs and injected with DHK ($\Delta F/F_0$ and frequency for baseline versus DHK $p<0.0001$; DHK versus CNO $\Delta F/F_0$ $p<0.0001$ and frequency $p<0.05$). **D)** Heat maps show $\Delta F/F_0$ calcium time courses (y-axis, 300 seconds scan time for each condition, >20% baseline) for all neurons (x-axis). Red line indicates each 300 seconds scan session, i.e. baseline, after injection of saline or DHK, and after subsequent injection of CNO.

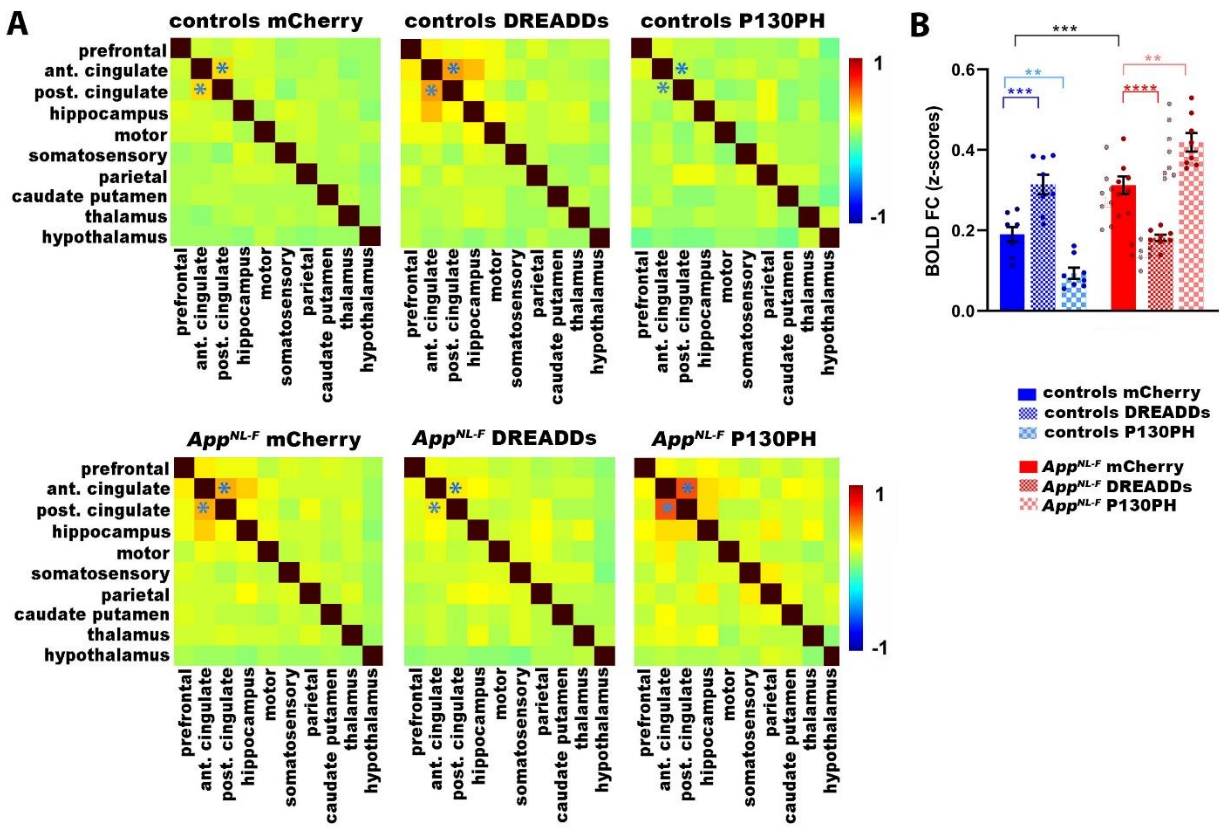


**<u>Fig. S6:</u> Exacerbation of astrocyte disruptions in *App*$^{NL\text{-}F}$ mice worsens hypersynchrony of BOLD signals in the cingulate cortex.** Astrocytes express either mCherry, DREADDs or P130PH (GFAP promoter) through local AAV injections in the cingulate cortex. **A)** BOLD FC matrices upon modulation of astrocyte calcium activity in control and *App*$^{NL\text{-}F}$ mice expressing mCherry, DREADDs or p130PH (N=8/group). FC between the anterior and posterior cingulate cortex is indicated by * on the matrices. **B)** Graphs show BOLD FC between the anterior and posterior cingulate cortex (z-scores ±SEM) for each group. **$p<0.01$, ***$p<0.001$, ****$p<0.0001$, one-way ANOVA with Sidak correction.

**Table S1: characteristics of F-PACK participants included for analysis.** For each characteristics the mean±SEM are shown per group.

| | **controls** | **Amyloid accumulators** |
|---|---|---|
| N | 24 | 10 |
| baseline CL | 7.1 ±1.1 | 7.7 ±2.1 |
| T1 CL | 6.1 ±1.6 | 42 ±5. |
| Age (years) | 78 ±1.0 | 77±2.0 |
| gender | N=12 female, N=12 male | N=5 female, N=5 male |
| baseline education (years) | 13 ±0.56 | 13 ±1.04 |
| baseline MMSE | 29 ±0.16 | 29 ±0.27 |

**Table S2: Seizure scoring**

| **score** | **Behaviour description** |
|---|---|
| 1 | Movement arrest |
| 2 | Neck jerks |
| 3 | Clonic seizures, sitting position |
| 4 | Myotonic seizures, flat on belly |
| 5 | Clonic or clonic-tonic seizures, sitting position |
| 6 | Clonic or clonic-tonic seizures with loss of balance, lying on side |

**References**


1. Saito T, et al. (2014) Single App knock-in mouse models of Alzheimer's disease. *Nat Neurosci* 17(5):661–663.

2. Roth BL (2016) DREADDs for Neuroscientists. *Neuron* 89(4):683–694.

3. Adamsky A, et al. (2018) Astrocytic Activation Generates De Novo Neuronal Potentiation and Memory Enhancement. *Cell* 174(1):59-71.e14.

4. Xie Y, Wang T, Sun GY, Ding S (2010) Specific disruption of astrocytic Ca2+ signaling pathway in vivo by adeno-associated viral transduction. *Neuroscience* 170(4):992–1003.

5. Goldey GJ, et al. (2014) Removable cranial windows for long-term imaging in awake mice. *Nat Protoc* 9(11). doi:10.1038/nprot.2014.165.

6. Shah D, et al. (2016) Early pathologic amyloid induces hypersynchrony of BOLD resting-state networks in transgenic mice and provides an early therapeutic window before amyloid plaque deposition. *Alzheimer's Dement* 12(9):964–976.

7. Grandjean J, Schroeter A, Batata I, Rudin M (2014) Optimization of anesthesia protocol for resting-state fMRI in mice based on differential effects of anesthetics on functional connectivity patterns. *Neuroimage* 102(P2). doi:10.1016/j.neuroimage.2014.08.043.

8. Cramer J V., et al. (2019) In vivo widefield calcium imaging of the mouse cortex for analysis of network connectivity in health and brain disease. *Neuroimage* 199. doi:10.1016/j.neuroimage.2019.06.014.

9. Batiuk MY, et al. (2020) Identification of region-specific astrocyte subtypes at single cell resolution. *Nat Commun* 11(1). doi:10.1038/s41467-019-14198-8.

10. Schaeverbeke JM, et al. (2021) Baseline cognition is the best predictor of 4-year cognitive change in cognitively intact older adults. *Alzheimer's Res Ther* 13(1). doi:10.1186/s13195-021-00798-4.

11. Adamczuk K, et al. (2013) Polymorphism of brain derived neurotrophic factor influences β amyloid load in cognitively intact apolipoprotein e ε4 carriers. *NeuroImage Clin* 2(1). doi:10.1016/j.nicl.2013.04.001.

12. Koole M, et al. (2009) Whole-body biodistribution and radiation dosimetry of 18F-GE067: A

radioligand for in vivo brain amyloid imaging. *J Nucl Med* 50(5). doi:10.2967/jnumed.108.060756.

13. Vandenberghe R, et al. (2010) 18F-flutemetamol amyloid imaging in Alzheimer disease and mild cognitive impairment a phase 2 trial. *Ann Neurol* 68(3). doi:10.1002/ana.22068.

14. Tzourio-Mazoyer N, et al. (2002) Automated anatomical labeling of activations in SPM using a macroscopic anatomical parcellation of the MNI MRI single-subject brain. *Neuroimage* 15(1). doi:10.1006/nimg.2001.0978.

15. Klunk WE, et al. (2015) The Centiloid project: Standardizing quantitative amyloid plaque estimation by PET. *Alzheimer's Dement* 11(1). doi:10.1016/j.jalz.2014.07.003.

16. De Meyer S, et al. (2020) Comparison of ELISA- and SIMOA-based quantification of plasma Aβ ratios for early detection of cerebral amyloidosis. *Alzheimer's Res Ther* 12(1). doi:10.1186/s13195-020-00728-w.

17. Reinartz M, et al. (2021) Changes in the language system as amyloid-β accumulates. *Brain*. doi:10.1093/brain/awab335.

18. Ran Q, et al. (2020) Reproducibility of graph measures at the subject level using resting-state fMRI. *Brain Behav* 10(8). doi:10.1002/brb3.1705.

19. Van Erum J, Van Dam D, De Deyn PP (2019) PTZ-induced seizures in mice require a revised Racine scale. *Epilepsy Behav* 95. doi:10.1016/j.yebeh.2019.02.029.
